# Lower-Limb Muscle Synergies in Musician’s Dystonia: A Case Study of a Drummer

**DOI:** 10.1101/2024.09.02.610624

**Authors:** Shizuka Sata, Kazuaki Honda, Satoshi Yamaguchi, Mizuki Komine, SungHyek Kim, Makio Kashino, Shota Hagio, Shinya Fujii

## Abstract

Musician’s dystonia (MD) is a movement disorder characterized by involuntary muscle contractions specifically triggered by playing an instrument. This condition often leads to a loss of fine motor control, threatening the careers of affected musicians. While MD is commonly associated with the hands, it can also affect the lower limbs, particularly in drummers. Understanding the muscle coordination involved in MD is crucial for comprehending its neurological mechanisms, yet the muscle coordination of lower-limb dystonia has not been thoroughly explored. This study aimed to investigate the differences in lower-limb muscle synergies in a drummer with MD, utilizing Non-negative Matrix Factorization (NMF) to analyze coordinated muscle activity patterns during drumming tasks. A 36-year-old male professional drummer with lower-limb MD was instructed to play a drum set in time with a metronome set at 80 beats per minute. The task involved striking the bass drum pedal in time with the downbeat. Electromyographic (EMG) data were collected from ten muscles in the right lower limb. The data were analyzed using NMF to extract muscle synergies and compare the number of synergies, spatial modules, and temporal modules between the data with and without dystonia symptoms. The number of muscle synergies did not differ significantly between the data with and without symptoms. Notably, changes were observed in both the spatial and temporal modules of muscle synergies. Spatial modules revealed the appearance of dystonia-specific muscle synergy, which is considered related to compensatory movement. Temporal modules showed significant earlier overactivation in timing, which is considered the direct manifestation of dystonia symptoms. These findings indicate that lower-limb dystonia in drummers affects the spatial and temporal profiles of muscle synergies. This study underscores the importance of considering both spatial and temporal modules of muscle synergy in understanding and treating lower-limb dystonia in drummers. Further research is needed to validate these findings and apply muscle synergy analysis for the clinical assessment of lower-limb dystonia in drummers.

## 1 Introduction

Focal task-specific dystonia (FTSD) is an involuntary movement disorder affecting some musicians due to maladaptive neuroplasticity (1-4). When this disorder manifests in musicians, it is referred to as musician’s dystonia (MD). It primarily induces involuntary movements specific to playing an instrument, presenting symptoms as a loss of fine motor control. This often leads to musicians abandoning their performance careers (5), making it a critical disorder for musicians.

While MD is most commonly reported in the hands, it also affects the lower limbs of drummers (6-8). In a case study of a drummer with lower limb MD, abnormal co-contraction in the thigh of the affected side during drum pedaling actions was noted (9). This drummer could alternately contract the plantar flexor and dorsiflexor muscles of the ankle on both affected and unaffected sides at a slow tempo. However, at increased tempos, he exhibited co-contraction of thigh muscles and was unable to maintain consistent performance at a fast tempo (9). In another case study of a drummer with lower limb MD, particularly in the right lower leg, it was revealed that the onset of symptoms led to increased activity in the ankle dorsiflexor muscles and partial thigh muscles, accompanied by a decrease in the activity of some toe extensor muscles (10). As a result, the amplitude of the played notes decreased, and synchronization errors increased (10). While these previous studies revealed symptom-specific activities in the individual lower limb muscles, how the symptom affected the coordination among the multiple muscles remains unclear.

By using a computational decomposition technique, such as Non-negative Matrix Factorization (NMF), it is possible to extract a small number of synchronized activation groups of muscles, referred to as muscle synergies, from the activities of multiple muscles (11-14). Previous studies have demonstrated that the activities of multiple muscles in cyclic movements, like locomotion, can be decomposed into several sets of weighting coefficients assigned to individual muscles, i.e., spatial modules, and activation coefficients related to the phase of movement, i.e., temporal modules (15-18). From the perspective of muscle synergies, there are at least three potential ways that muscle coordination could be altered when dystonia symptoms manifest. First, the number of muscle synergies could differ. A reduction or increment in the number of muscle synergies suggests affecting the number of motor modules that can be independently recruited (19, 20). Second, the spatial modules of muscle synergy could differ. Changes in the spatial modules would indicate that dystonia primarily affects the extent to which each muscle participates in movement. Third, the temporal modules of muscle synergy could vary. Changes in temporal modules would show that the dystonia impacts the time-dependent profiles of when and how strongly each muscle synergy is activated to perform the movement. Together, investigating how dystonia symptoms influence 1) the number of synergies, 2) spatial modules, and 3) temporal modules is crucial. Clarifying these aspects would contribute to understanding how FTSD affects muscle coordination and elucidate clinical outcomes and neural underpinnings.

A central question of this study is how muscle synergies differ when dystonic symptoms occur in a drummer with lower limb dystonia. The previous research on pianists with MD revealed that while the number of muscle synergies remained unchanged, partial changes were observed in the coordination structures (spatial modules) between the affected and unaffected hands, as well as in comparison to the hands of healthy pianists (21). Moreover, these changes were identified as either directly related to the dystonic symptoms or as compensatory, based on their association with performance accuracy (21). This suggests that occurrence of dystonia symptoms affects the spatial modules without changes in the number of synergies. Another previous research in childhood dystonia reported the differences in the temporal modules while the number of synergies remained unchanged (22). This suggests that occurrence of dystonia symptoms affects the temporal modules without changes in the number of synergies. Taken together, the previous studies on muscle synergies in FTSD suggest changes in either spatial or temporal modules, but not in the number of synergies. Considering these previous studies on muscle synergies in FTSD, we hypothesized the following: 1) The drummer with lower-limb dystonia would show no significant change in the number of muscle synergies, while there would be changes in 2) the spatial modules, and/or 3) the temporal modules. The aim of this study was to investigate how the number of muscle synergies and the spatial and temporal modules would differ when dystonic symptoms occur in a drummer with lower-limb dystonia and to test the hypotheses in a drummer with right lower-limb dystonia while performing a drum pattern.

## 2 Methods

### Participant

The same participant as Honda et al. participated in this study (10). The participant was a 36-year-old male professional rock drummer. He began playing drums at age 14 and was diagnosed by a neurologist with focal task-specific dystonia in his right lower limb at age 29. He first experienced symptoms at the age of 24 while on a national tour. He complained that the right lower leg involuntarily contracted during drum playing. At times when his symptoms were most severe, he felt discomfort in his right foot during activities that resembled drum pedaling motions, such as ascending stairs and driving a vehicle. To mitigate these symptoms, he utilized sensory tricks, such as adjusting the height of his shoes and chair, which provided only temporary relief. Eventually, the progression of focal task-specific dystonia impaired his ability to play the drums, leading to his withdrawal from public performances. His family history revealed no neurological disorders. His gait was normal, and he had no other neurological diseases. He was prescribed no medication for at least the past 3 years. He had no history of other neuropsychiatric disorders or neurosurgery. Ethical approval for this study was obtained from the Communication Science Laboratories Research Ethics Committee at Nippon Telegraph and Telephone Corporation (Approval Number: H30-009). The experiment was conducted according to principles originating in the Declaration of Helsinki. Written informed consent was obtained from the participant in this study.

### Experimental task

The participant was instructed to play an eight-beat drum pattern on a drum set in time with a metronome sound set at a constant tempo of 80 beats per minute (bpm). The score of the drum pattern is depicted in Figure 1. The drumming pattern consisted of a single chunk of four metronome sounds that indicated the beat position in the pattern (see the vertical arrows in Figure 1), defined as 1 bar. Each trial consisted of 60 bars, and the participant completed a total of four trials. The affected right lower limb was used to play the bass drum, striking the drum pedal on the downbeat of the first beat and the syncopated upbeat of the third beat. The downbeat corresponds to the beginning of a beat, while the upbeat is situated between consecutive downbeats, such as when the participant played between the third and fourth beats. At the beginning of each trial, the participant started playing after hearing a 1 kHz pure tone and four metronome tones, which served as a cue for the start of the trial and to signal the tempo. The cue signal mimicked the standard practice in live performances, where a four-beat count-in is used. The participant was allowed a rest period of at least 1 minute between trials. During the drum pattern, the participant verbally reported the occurrence of symptoms whenever he felt an abnormality in his movements.

**Figure 1.**
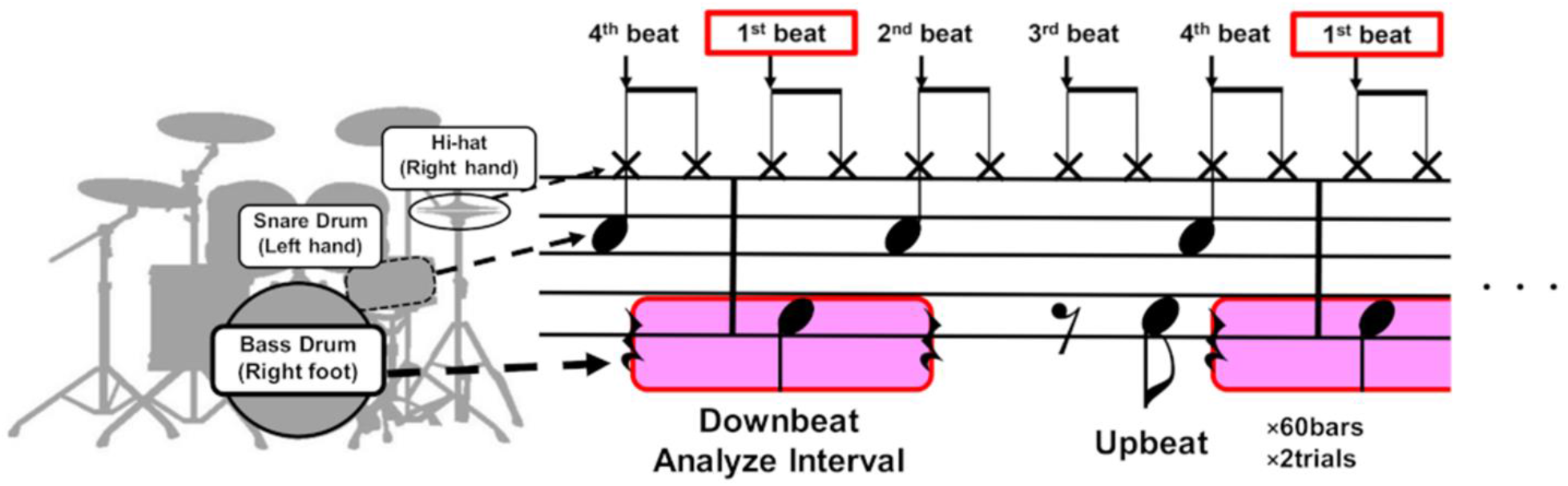
The score of the drum pattern used in the experiment. The participant was instructed to play an eight-beat drum pattern on a drum set in time with a metronome consisting of a single chunk of four metronome sounds that indicated the beat position in the pattern (see the vertical black arrows). The drumming pattern involved playing the bass drum with the right foot, the snare drum with the left hand, and the hi-hat cymbals with the right hand. The participant mostly exhibited dystonia symptoms in the right lower limb when playing the downbeat of the first beat (see the areas highlighted in pink). We therefore analyzed the muscle synergy of the right lower limb before and after the downbeat timings.

### Data collection

Electromyographic (EMG) activities were measured using active bipolar Ag/AgCl surface electrodes with a Trigno Wireless EMG system (DELSYS Corp., Boston, MA, USA). The EMG activities were recorded from ten muscles in the right lower extremity: rectus femoris (RF), lateral head of biceps femoris (BF), vastus lateralis (VL), vastus medialis (VM), tibialis anterior (TA), extensor digitorum longus (EDL), gastrocnemius (GAS), soleus (SOL), peroneus longus (PL), and extensor digitorum brevis (EDB), at a sampling rate of 1,111 Hz. The interelectrode distance was 10 mm. A trigger signal from the EMG system was sent to an audio interface (Fireface UCX: RME Corp., Germany), and the metronome sounds were recorded synchronously with the EMG signals at a sampling rate of 48,000 Hz.

### Data preprocessing

Of the four trials, the first trial was excluded due to the failure of EMG recording, and the fourth trial was excluded from the analysis because the participant self-reported that he intentionally changed his drumming movements compared to the other trials. Consequently, we analyzed the data from the second and third trials to investigate muscle synergy. Since each of the trial consisted of 60 bars, there were 120 bars of the data in the second and third trials. During the second trial, the participant verbally reported 11 occurrences of symptoms on the downbeat of the first beat and 2 occurrences on the upbeat of the third beat. In the third trial, the participant reported 9 occurrences of symptoms on the downbeat of the first beat but no occurrences of symptoms on the upbeat of the third beat. Taken together, the participant reported 20 occurrences of symptoms on the downbeat of the first beat and 2 occurrences on the upbeat of the third beat in the second and third trials. Because the participants reported the symptoms mostly on the downbeat of the first beat, we decided to analyze the right lower-limb EMG data of playing the bass drum for the first beat. In total, there were 20 beats of data with dystonia and 100 beats of data without dystonia. Since three of the EMG data without dystonia included artifacts caused by intense electrode vibration during body movement, we excluded these data from the analysis. Thus, we used 20 beats of EMG data with dystonia and 97 beats of EMG data without dystonia for muscle synergy analysis.

The EMG signals were demeaned, full-wave rectified, and low-pass filtered using the 4th order Butterworth filter with a cutoff of 20Hz. The metronome sound signals were rectified, and the envelope was calculated to detect the peak timings, which were used as reference timings. We extracted the EMG signals 0.75 seconds before and after the reference timings. The interval of 0.75 seconds was selected because the metronome tempo was set at a constant tempo of 80 bpm, resulting in intervals of approximately 0.75 seconds between the reference timings (i.e., 60 seconds / 80 beats = 0.75 seconds / beat). Each of the extracted EMG data within this interval included the lifting and lowering of lower-limb movements to kick the bass drum in synchronization with the metronome sounds.

Each of the extracted EMG data over the 1.50 sec, termed as one cycle of EMG data, was resampled into 151 time points. Since we recorded the EMG from 10 muscles, we formed an EMG data matrix consisting of 10 muscles × 20 cycles for the data with dystonia, and an EMG data matrix consisting of 10 muscles × 97 cycles for the data without dystonia. The minimum and maximum values across the cycles were calculated for each of the 10 muscles and normalization was performed for each muscle vector by subtracting the minimum value and dividing it by the maximum value. Thus, each muscle vector in the EMG data matrix was divided by standard deviation of each muscle vector, *σ*, to have unit variance, ensuring that the activity of each muscle was equally weighted (15).

### Overview of muscle synergy analysis

Muscle synergy analyses were performed in two steps to examine the commonalities and differences between the EMG activities with and without dystonia symptoms. In the first step, muscle synergies were extracted for each of the EMG activities with and without dystonia. This first step aimed to determine the number of muscle synergies in each of the dataset. In the second step, we pooled the EMG data with and without dystonia, and muscle synergies were simultaneously extracted from the pooled dataset. The second step aimed to identify the shared and specific muscle synergies (23).

Specifically, we aimed to extract the commonalities and differences in the muscle synergy due to the presence or absence of dystonia symptoms.

### Muscle synergy analysis for each of the EMG data with and without symptoms

Muscle synergies were extracted for each of the EMG data matrices with and without dystonia symptoms using NMF (11, 14, 24). NMF assumes that a given muscle activation pattern *M* at each point in time is composed of a linear combination of several muscle weight vectors *W*_*i*_, each recruited by activation coefficients *C*_*i*_ (11, 14, 24). Therefore, a specific muscle activation pattern *M* can be expressed as follows:

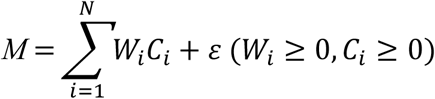

where *i* represents the relative contribution of muscles involved in the synergy, *N* denotes the number of synergies, and *ε* represents the residual. To compare muscle weight vectors of synergies between the data with and without symptoms with the same scaling as the measured EMG activity, a unit variance scaling was reverted by multiplying each weight of the *i*^*th*^ muscle by *σ*_*i*_ for all muscle synergies. The muscle weight and activation coefficient matrices were normalized so that each weight vector became a unit vector.

We first tested whether the number of muscle synergies was similar between the data with and without dystonia. For the EMG dataset with dystonia, the data across 20 cycles were randomly split into an 80% training set (i.e., 16 cycles) and a 20% test set (i.e., 4 cycles) for cross validation (25, 26). The test EMG datasets were reconstructed by muscle-weighting matrices derived from 1 to 10 muscle synergies extracted from the training EMG datasets. This cross-validation procedure was repeated 20 times. Then, the goodness-of-fit of the data reconstruction was quantified for each number of muscle synergies by the average value of the variance accounted for (VAF), which was defined as a 100 × the uncentered Pearson correlation coefficient (27, 28). For the EMG dataset without dystonia, 20 cycles of data were randomly selected from the 97 cycles. The cross-validation procedure was the same as for the data with dystonia but repeated 1,000 times to estimate the 95% bootstrapping confidence interval (95% CI) of the VAF value. This series of procedures aimed to confirm whether the VAF value from the EMG dataset with symptoms fell within the range of inter-cycle variability of the EMG dataset without dystonia. To determine the number of muscle synergies, a least squares method was used to fit a line to the portion of the VAF curve, identifying the point at which the VAF curve linearly plateaued as the number of muscle synergies where the mean squared error (MSE) falls below 10^−5^ (26).

For the EMG dataset without symptoms, to estimate the 95% CI of the muscle weighting vectors and activation coefficients of muscle synergies, the sets of muscle synergies were extracted 1,000 times across randomly selected EMG datasets each consisting of 20 cycles. Using k-means clustering, the entire set of muscle synergies were classified into clusters based on the cosine of the angle between pair of muscle weight vectors (15). K-means clustering was applied with pairwise constraints, preventing muscle synergies calculated in any of the 1,000 iterations from being classified into the same cluster (15, 29). The number of clusters was defined as the same determined based on the VAF curve as described above.

### Shared and specific muscle synergy analysis

The EMG dataset with symptoms (20 cycles) and the EMG dataset without dystonia (97 cycles) were combined into a single EMG data matrix to extract muscle synergies that explain the entire EMG data matrix (i.e., shared synergies) and each of the datasets with and without dystonia (i.e., specific synergies) (23). First, the total number of muscle synergies to be extracted was set to the sum of the number of muscle synergies determined for each of the datasets with and without dystonia, assuming that there was no muscle synergies shared between the datasets with and without dystonia.

Consequently, the activation coefficients specific to the dataset with dystonia were set to zero for the muscle synergies without dystonia, and vice versa. The total number of extracted muscle synergies was then reduced one by one in each extraction procedure, and the number of shared and specific muscle synergies was adjusted. The number of shared and specific muscle synergies was determined as the minimum number of muscle synergies required to exceed the VAF value extracted in the analyses for each of the data with and without dystonia.

## 3 Results

### The number of muscle synergies

The VAF curves linearly plateaued as the number of muscle synergies for each of the datasets with and without dystonia symptoms (see red and blue lines, respectively, in Figure 2). For the data with symptoms (red), the minimum number of muscle synergies for which the MSE calculated from the VAF curve was below 10^−5^ was 6 (MSE = 7.22.10^−6^). The 6 muscle synergies explained 96.46% of the original EMG data variance with dystonia. For the data without dystonia (blue), the minimum number of muscle synergies for which the MSE calculated from the VAF curve was below 10^−5^ was also 6 (MSE = 6.79.10^−6^). The 6 muscle synergies explained 92.36-96.99% (lower and upper limits of 95% CIs) of the original EMG data variance without dystonia. The VAF values with 1 to 7 muscle synergies to account for the EMG data with dystonia were within the 95% CI range for those without dystonia, indicating that the number of synergies was the same to account for both the data with and without dystonia.

**Figure 2.**
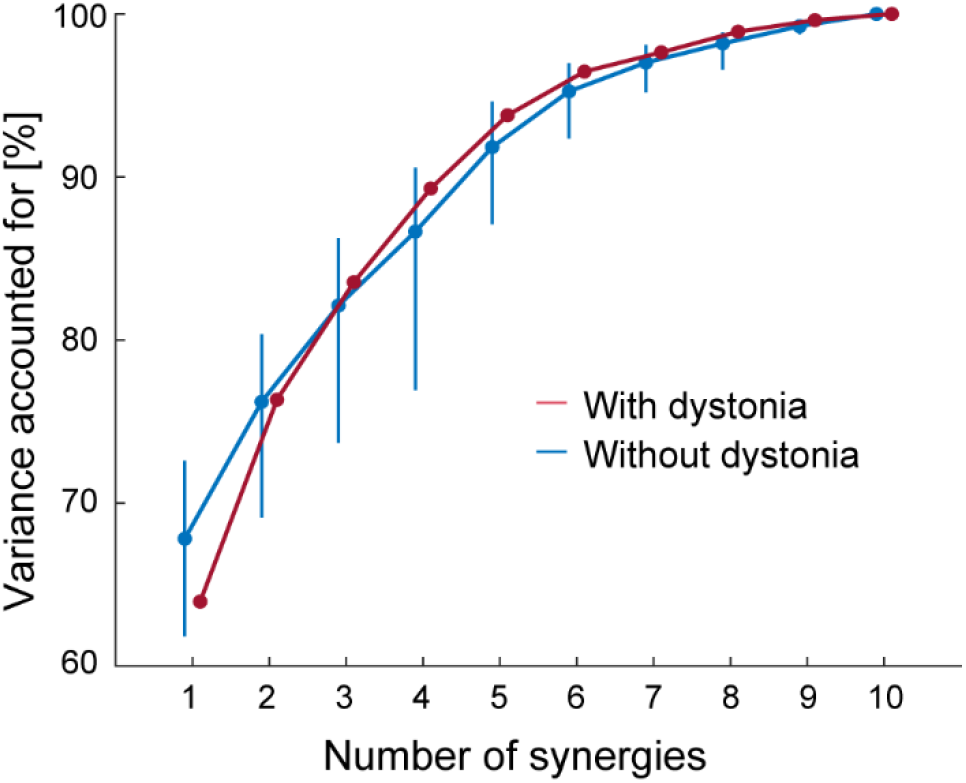
The variance accounted for (VAF) as a function of the number of muscle synergies. The VAF for the EMG data with dystonia symptoms is shown in red, and that for the data without dystonia symptoms is shown in blue. The blue bars represent the upper and lower limits of the 95% bootstrapping confidence intervals (CIs) of the VAF for each number of muscle synergies in the data without dystonia symptoms.

### Muscle synergies for each of the data with and without dystonia

The 6 muscle synergies extracted for each of the datasets with and without dystonia symptoms are shown in Figure 3. The spatial modules of muscle synergy, or the extracted muscle-weight vectors, are shown in the left panels of Figure 3, while the temporal modules of muscle synergy, or the extracted time-dependent profiles of when and how strongly each muscle synergy was activated, are shown in the right panels of Figure 3.

**Figure 3.**
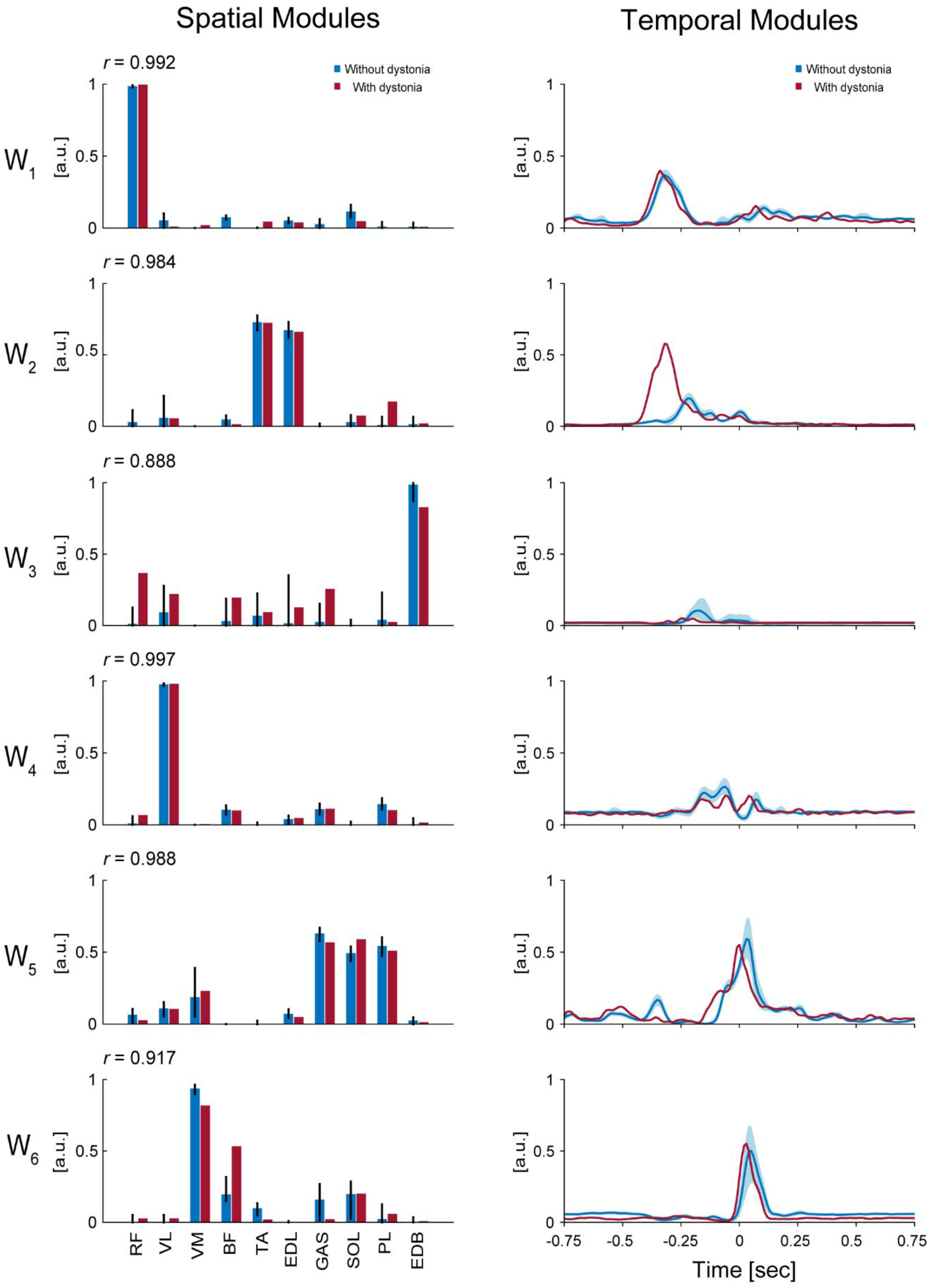
Muscle synergies for each of the data with and without dystonia. The spatial and temporal modules of muscle synergies are shown in the left and right panels, respectively. The red bars and lines indicate the spatial and temporal modules extracted from the data with dystonia, while the blue bars and lines indicate those from the data without dystonia. In the left panels, the bars represent muscle weights of rectus femoris (RF), vastus lateralis (VL), vastus medialis (VM), biceps femoris (BF), tibialis anterior (TA), extensor digitorum longus (EDL), soleus (SOL), gastrocnemius (GAS), peroneus longus (PL), and extensor digitorum brevis (EDB), respectively, extracted from the non-negative matrix factorization (NMF). The error bars represent the 95% CI in the data without dystonia. The *r* value indicates the cosine similarity between the muscle-weight vectors with and without dystonia. In the right panels, the activation coefficients in the NMF, or the temporal modules of muscle synergy, are shown 0.75 seconds before and after the metronome sound. The order of the muscle synergies (W_1_-W_6_) was sorted by the time of peak activation of the temporal modules.

The first spatial module, or the extracted muscle-weight vector (W_1_), showed a high weighting for the RF muscle, which is responsible for hip flexion. The cosine similarity demonstrated the commonality in the use of this spatial module between the data with and without symptoms (*r* = 0.992). The time-dependent profiles (i.e., temporal modules) of this muscle synergy showed peak values at approximately 338 milliseconds (ms) and 318 ms before the metronome sound for the data with and without dystonia, respectively. The peak values were 0.40 [arbitrary units (a.u.)] and 0.37 [a.u.] for the data with and without dystonia, respectively. Both the spatial and temporal modules of this muscle synergy were similar between the data with and without dystonia, indicating that W_1_ for hip flexion to lift the thigh in preparation for the kicking movement was used in a similar manner, irrespective of the presence or absence of symptoms.

The second spatial module (W_2_) showed high weightings for the TA and EDL muscles, which are responsible for ankle dorsiflexion. The cosine similarity demonstrated the commonality in the use of this spatial module between the data with and without dystonia (*r* = 0.984). However, the temporal modules of this muscle synergy differed between the data with and without dystonia. Specifically, the temporal module shifted earlier and was more activated in the data with symptoms compared to those without dystonia: the peak values were observed at approximately 318 ms and 219 ms before the metronome sound for the data with and without dystonia, respectively. The peak values were 0.58 [a.u.] and 0.20 [a.u.] for the data with and without dystonia, respectively. Thus, while the spatial modules were similar, the temporal modules differed between the data with and without dystonia, indicating that the temporal module of ankle dorsiflexion muscle synergy to prepare for the kicking movement occurred earlier and over-activated when the symptoms occurred.

The third spatial module (W_3_) showed a relatively high weighting for the EDB muscle, which is responsible for toe extension, and relatively low weightings for the other muscles (RF, VL, BF, TA, EDL, GAS, and PL). The cosine similarity of W_3_ demonstrated relatively lower values compared to the other spatial modules (*r* = 0.868). The temporal modules of this muscle synergy also differed between the data with and without dystonia. Specifically, the temporal module shifted earlier and was less activated in the data with dystonia compared to those without dystonia: the peak values were observed at approximately 249 ms and 179 ms before the metronome sound for the data with and without dystonia, respectively. The peak values were 0.05 [a.u.] and 0.11 [a.u.] for the data with and without dystonia, respectively. Thus, both the spatial and temporal modules differed between the data with and without dystonia.

The fourth spatial module (W_4_) showed a high weighting for the VL muscle, which is responsible for stabilizing the hip joint. The cosine similarity demonstrated the commonality in the use of this spatial module between the data with and without dystonia (*r* = 0.997). The temporal modules of this muscle synergy showed peak values at approximately 60 ms before the metronome sound for both the data with and without dystonia. The peak values were 0.20 [a.u.] and 0.26 [a.u.] for the data with and without dystonia, respectively. Nevertheless, the temporal modules of this muscle synergy were slightly different between the data with and without dystonia.

The fifth spatial module (W_5_) showed high weightings for the GAS, SOL, and PL muscles, which are responsible for plantar flexion. The cosine similarity demonstrated the commonality in the use of this spatial module between the data with and without dystonia (*r* = 0.991). The temporal modules of this muscle synergy showed peak values around the metronome sound for the data with symptoms and approximately 30 ms after the metronome sound for the data without dystonia. The peak values were 0.55 [a.u.] and 0.59 [a.u.] for the data with and without dystonia, respectively. The shape of the temporal modules of this muscle synergy was overall similar but appeared slightly shifted earlier in the data with dystonia compared to those without dystonia.

The sixth spatial module (W_6_) showed relatively high weightings for the VM and BF muscles, which are responsible for stabilizing the knee and hip joints. The cosine similarity of W_6_ was *r* = 0.922, which was relatively lower compared to the other spatial modules. Specifically, the weighting for the VM was lower, but the weighting for the BF was higher in the data with dystonia. The temporal modules of this muscle synergy showed peak values at approximately 30 ms and 60 ms after the metronome sound for the data with and without dystonia, respectively. The peak values were 0.55 [a.u.] and 0.50 [a.u.] for the data with and without dystonia, respectively. The shape of the temporal modules of this muscle synergy was overall similar but appeared slightly shifted earlier in the data with dystonia compared to those without dystonia.

### Shared and specific muscle synergies

The shared and specific muscle synergy analysis showed a VAF of 95.79% for the combined dataset, a VAF of 95.45% for the data with dystonia, and a VAF of 94.96% for the data without dystonia. The analysis revealed that there were 5 shared muscle synergies between the data with and without dystonia symptoms, while there were 2 specific muscle synergies each for the data with and without dystonia symptoms (Figure 4).

**Figure 4.**
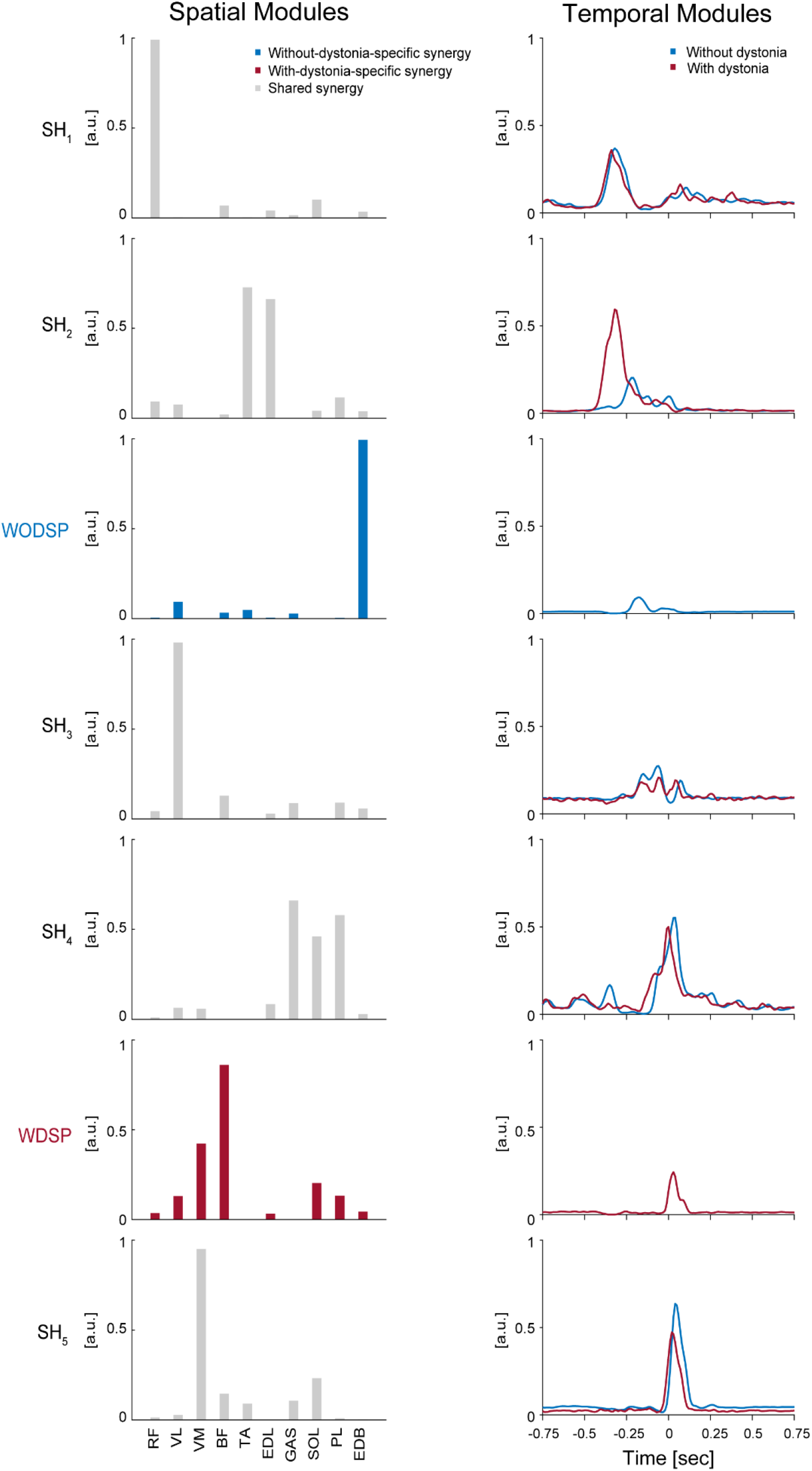
Shared muscle synergies between the data with and without dystonia and specific muscle synergies for each dataset. The spatial and temporal modules of shared and specific muscle synergies are shown in the left and right panels, respectively. In the left panels, the bars represent muscle weights of rectus femoris (RF), vastus lateralis (VL), vastus medialis (VM), biceps femoris (BF), tibialis anterior (TA), extensor digitorum longus (EDL), soleus (SOL), gastrocnemius (GAS), peroneus longus (PL), and extensor digitorum brevis (EDB), respectively, extracted from the non-negative matrix factorization (NMF).The grey bars represent the shared spatial modules between the data with and without dystonia (SH_1-5_). The blue bars represent the without-dystonia-specific synergy (WODSP). The red bars represent the with-dystonia-specific synergy (WDSP). In the right panels, the activation coefficients in the NMF, or the temporal modules of muscle synergy, are shown 0.75 seconds before and after the metronome sound. The red lines indicate the temporal modules for the data with dystonia, while the blue lines indicate those for the data without dystonia.

The first shared spatial module (SH_1_) showed a high weighting for the RF muscle, which is responsible for hip flexion. The time-dependent profiles (i.e., temporal modules) of this muscle synergy were similar between the data with and without dystonia, indicating that SH_1_ for hip flexion to lift the thigh in preparation for the kicking movement was shared between the data with and without dystonia symptoms.

The second shared spatial module (SH_2_) showed high weightings for the TA and EDL muscles, which are responsible for ankle dorsiflexion. The temporal modules of this muscle synergy clearly differed between the data with and without dystonia. Specifically, the temporal module shifted earlier and was more activated in the data with dystonia compared to those without symptoms. Thus, this shared spatial module was activated more and earlier when symptoms occurred.

The without-dystonia-specific synergy (WODSP) showed a high weighting for the EDB muscle, which is responsible for toe extension (see blue in Figure 4). This muscle synergy appeared specifically in the data without dystonia, indicating that the muscle synergy for toe extension during the preparation movement was observed when dystonia symptoms did not occur.

The third shared spatial module (SH_3_) showed a high weighting for the VL muscle, which is responsible for stabilizing the hip joint. The shape of the temporal modules of this muscle synergy was overall similar between the data with and without dystonia but appeared slightly shifted earlier in the data with dystonia compared to those without dystonia.

The fourth shared spatial module (SH_4_) showed high weightings for the GAS, SOL, and PL muscles, which are responsible for plantar flexion. The shape of the temporal modules of this muscle synergy was overall similar between the data with and without dystonia but appeared slightly shifted earlier in the data with dystonia compared to those without dystonia.

The with-dystonia-specific synergy (WDSP) showed relatively high weightings for the BF and VM muscles, and relatively low weightings for the RF, VL, EDL, SOL, PL, and EDB muscles. The temporal module showed activation shortly after the metronome sound, indicating that this muscle synergy was contributing to stabilizing the kick movement after pedaling specifically when the dystonia symptoms occurred.

The fifth shared spatial module (SH_5_) showed a high weighting for the VM muscle. The shape of the temporal modules of this muscle synergy was overall similar between the data with and without dystonia but appeared slightly shifted earlier in the data with dystonia compared to those without dystonia.

## 4 Discussion

By utilizing NMF to extract the spatial modules (i.e., muscle-weight vectors) and the temporal modules (i.e., activation coefficients), we aimed to investigate how the number of muscle synergies and the spatial and temporal modules would differ when dystonic symptoms occur in a drummer with lower-limb dystonia. Specifically, we tested whether 1) the drummer with lower-limb dystonia would show no significant change in the number of muscle synergies, while there would be changes in 2) the spatial modules, and/or 3) the temporal modules when the dystonia symptoms occurred. Our results showed that, while the number of muscle synergies was the same in accounting for both the data with and without dystonia, the spatial and temporal modules of the muscle synergies changed when the dystonia symptoms occurred.

### Number of synergies

Our first hypothesis was that the drummer with lower-limb dystonia would show no significant change in the number of muscle synergies when the dystonia symptoms occurred. Our results showed that 6 muscle synergies explained 96.46% and 92.36-96.99% of the original EMG data variance in the data with and without dystonia, respectively (Figure 2). These results indicate that six muscle synergies account for both the data with and without symptoms, and the number of synergies was the same regardless of the occurrence of symptoms. Thus, our results support the first hypothesis and align with previous studies on MD pianists (21) and dystonia in children (22), suggesting that the number of synergies or motor modules that can be independently recruited remains unchanged when dystonia symptoms occur.

Nevertheless, it is worth mentioning that previous studies on patients with stroke (19, 30), cerebral palsy (31), spinal cord injury (32), and Parkinson’s disease (33) have shown changes in the number of muscle synergies. Specifically, these studies reported a reduction in the number of muscle synergies due to these neurological disorders. The reduced number of synergies indicates that motor commands become simplified and less complex when these neurological disorders occur. In contrast, we did not observe a reduction in the number of synergies, suggesting a specific nature of FTSD compared to other neurological conditions such as stroke, cerebral palsy, spinal cord injury, and Parkinson’s disease. Our results suggest that the lower-limb dystonia in the drummer cannot be explained by simplified motor commands or a reduced number of muscle synergies.

### Spatial modules

Our second hypothesis was that there would be changes in the spatial modules of muscle synergies when the dystonia symptoms occurred. Our results showed that the structure of the spatial modules differed when the dystonia symptoms occurred. Specifically, the shared and specific muscle synergy analyses revealed that there were 5 shared muscle synergies between the data with and without dystonia symptoms, while there were 1 specific muscle synergies each for the data with and without dystonia symptoms (Figure 4). These results support the second hypothesis, indicating that the spatial modules of muscle synergies change when symptoms occur in the drummer with lower-limb dystonia.

It is noteworthy that we observed the without-dystonia-specific synergy (WODSP), which had a high weighting for the EDB muscle (blue in Figure 4). Since this muscle synergy was activated before the kicking movement and the EDB muscle contributes to toe extension, this synergy seemed to contribute to the fine control of toe extension in preparation for the kicking movement when the dystonia symptoms did not occur. In contrast, when the dystonia symptoms occurred, we observed the with-dystonia-specific synergy (WDSP), which had high weightings for the BF and VM muscles (red in Figure 4). Since this muscle synergy was activated after the metronome sound, it appeared to contribute to stabilize the lower limb after the kicking movement. Thus, although the number of synergies was the same—six (5 shared plus 1 specific synergy)—between the data with and without dystonia, we found a different structure of the spatial modules depending on the absence or occurrence of the symptoms.

The previous study on pianists with MD showed changes in the spatial modules of muscle synergies between the affected and unaffected hands, as well as in comparison to the hands of healthy pianists (21). These changes in the spatial modules were identified as either directly related to the dystonic symptoms or as compensatory (21). We suggest that the with-dystonia-specific synergy (WDSP) observed in this study may be interpreted as compensatory rather than as a direct manifestation of dystonic symptoms. This interpretation is based on the fact that the WDSP was observed relatively late in the kicking movement. The spatial module specific to dystonia symptoms in this study may therefore be understood as a muscle synergy that compensates for the movement after the direct manifestation of dystonic symptoms.

### Temporal modules

What kind of muscle synergy would reflect the direct manifestation of symptoms in the drummer with lower-limb dystonia? Based on the results of the temporal modules in this study, we propose that a change in the temporal module of muscle synergy reflects the direct manifestation of dystonic symptoms in the drummer. Specifically, we found that the temporal modules of the second shared spatial module (SH_2_), which had high weightings for the TA and EDL muscles, clearly differed between the data with and without symptoms: the temporal module shifted earlier and was more activated in the data with dystonia compared to those without dystonia (Figure 4). This synergy was activated at almost the same timing as the first shared spatial module (SH_1_), suggesting that the temporal module intended to activate SH_1_ may have contaminated the other muscle synergy and driven SH_2_ when the symptoms occurred.

What could be the neural mechanisms related to the earlier-shifted overactivity of the muscle synergy (SH_2_) observed in this study? We suggest that abnormal inhibition and/or excitation of motor commands driving the muscle synergy may be one of the mechanisms. Previous neuroimaging studies have shown abnormal overactivity of the primary motor cortex (M1) in patients with FTSD (34, 35). Transcranial magnetic stimulation (TMS) studies have also shown a loss of inhibition in M1 in patients with FTSD (36,37,38). Moreover, a recent study on pianists with FTSD showed reduced inhibition and elevated facilitation in M1 compared to healthy controls (39). Considering these previous studies, we assume that the overactivity or loss of inhibition in M1 might be related to the earlier-shifted overactivity of the muscle synergy SH_2_. It might be possible that the motor commands driving the muscle synergy for hip flexion (SH_1_) contaminated the muscle synergy for ankle dorsiflexion (SH_2_) due to the overactivity or loss of inhibition in M1.

We suggest that the occurrence of the earlier-shifted overactivity of the muscle synergy SH_2_ concealed the without-dystonia-specific synergy (WODSP) and induced slight shifts in the temporal-module activities of SH_3_, SH_4_, and SH_5_. This, in turn, might have resulted in the appearance of the with-dystonia-specific synergy (WDSP) to compensate for the movement. Specifically, the enhanced dorsiflexion movement due to the earlier-shifted overactivity of the muscle synergy SH_2_ during the pedaling preparation phase might lead to a decrease in the force applied to the pedal. To compensate for this reduced force, the with-dystonia-specific synergy (WDSP) might appear to provide the necessary force for pedaling despite the disrupted motor control caused by the dystonia symptoms.

In a previous study of lower-limb dystonia in drummers by Honda et al., it was reported that the appearance of dystonia symptoms resulted in the earlier timing of bass drum performance (10). We suggest that the earlier shift of temporal-module activity of SH_2_, together with the slight shifts in those of SH_3_, SH_4_, and SH_5_, caused the earlier timing of bass drum performance. The without-dystonia-specific synergy (WODSP) is considered to contribute to the fine control of toe extension in preparation for the kicking movement. This might help achieve a “whip-like motion” or the sequential proximal-to-distal motion in preparation for the kicking movement. Similar preparatory movements have been reported in skilled pianists, where a lifting of the upper limb from the proximal to the distal end precedes the keystroke (40), suggesting a common motor strategy among musicians to facilitate skilled performance. We suggest that the earlier-shifted overactivity of the muscle synergy SH_2_ hindered the “whip-like” preparation for the kicking movement and resulted in compensatory movement caused by the with-dystonia-specific synergy (WDSP). Taken together, we propose that the change in the temporal module of muscle synergy reflects the direct manifestation of dystonic symptoms in the drummer with lower-limb dystonia, and this supports our third hypothesis.

### Clinical implications and limitations

As far as we know, this is the first case study to apply NMF to investigate the muscle synergy of the lower limb in a drummer with FTSD. So far, there have been a limited number of studies on drummers with FTSD (6-10). Therefore, formal assessments and clinical interventions for lower limb dystonia in drummers have not been fully explored. Based on the results of this study, we suggest that NMF and muscle synergy analysis may be useful tools for assessing the symptoms of lower-limb dystonia. Specifically, monitoring the spatial and temporal modules of muscle synergies could be beneficial for symptom assessment. However, given the limited research on drummer’s dystonia, more studies are needed before clinical applications can be developed. For instance, future research should expand the sample size of MD drummers and compare them to healthy drummers to validate the findings from this study. Longitudinal studies tracking changes in muscle synergies over time with various interventions would also be valuable in understanding the progression and potential recovery mechanisms of dystonia in musicians.

## 5 Conclusion

By applying NMF to 10 lower-limb muscles in a drummer with FTSD, we found that the number of muscle synergies did not differ between the data with and without dystonia; however, changes were observed in both the spatial and temporal modules of muscle synergies due to the appearance of symptoms. Spatial modules revealed the appearance of dystonia-specific muscle synergy, which is considered related to compensatory movement. Temporal modules showed significant earlier overactivation in timing, which is considered the direct manifestation of dystonia symptoms. These findings indicate that lower-limb dystonia in drummers affects the spatial and temporal profiles of muscle synergies, and NMF and muscle synergy analysis may be useful tools for assessing the symptoms of a drummer’s lower-limb dystonia.

## 6 Conflict of Interest

Kazuaki Honda and Makio Kashino are employed by the NTT Communication Science Laboratories, Nippon Telegraph and Telephone Corporation, Japan. The remaining author has no relevant financial or non-financial interests to disclose. There is no product in development or marketed products to declare.

## 7 Author Contributions

SS: Data curation, Formal Analysis, Funding acquisition, Investigation, Methodology, Validation, Visualization, Writing original draft, Writing – review & editing. KH: Data curation, Formal Analysis, Methodology, Software, Writing – review & editing. SY: Resources, Writing – review & editing. MiK: Data curation, Writing – review & editing. SK: Methodology, Writing – review & editing. MaK: Resources, Writing – review & editing. SH: Conceptualization, Formal Analysis, Investigation, Methodology, Supervision, Validation, Visualization, Writing – original draft, Writing – review & editing. SF: Conceptualization, Funding acquisition, Investigation, Methodology, Project administration, Supervision, Validation, Visualization, Writing original draft, Writing – review & editing.

## 8 Funding

This work was supported by Taikichiro Mori Memorial Research Grant, and JST SPRING Grant (no. JPMJSP2123) (to S.S.); by a JST COI-NEXT grant (no. JPMJPF2203), a JST PRESTO grant (no. JPMJPR23S9), and a KGRI Pre-startup Grant (to S.F.).

